# PUFA stabilizes a conductive state of the selectivity filter in IKs channels

**DOI:** 10.1101/2024.01.11.575247

**Authors:** Alessia Golluscio, Jodene Eldstrom, Jessica J. Jowais, Marta E. Perez-Rodriguez, Kevin P. Cunningham, Alicia de la Cruz, Xiaoan Wu, Valentina Corradi, D. Peter Tieleman, David Fedida, H. Peter Larsson

## Abstract

In cardiomyocytes, the KCNQ1/KCNE1 channel complex mediates the slow delayed-rectifier current (IKs), pivotal during the repolarization phase of the ventricular action potential. Mutations in IKs cause Long QT Syndrome (LQTS), a syndrome with a prolonged QT interval on the ECG, which increases the risk of ventricular arrhythmia and sudden cardiac death. One potential therapeutical intervention for LQTS is based on targeting IKs channels to restore channel function and/or the physiological QT interval. Polyunsaturated fatty acids (PUFAs) are potent activators of KCNQ1 channels and activate IKs channels by binding to two different sites, one in the voltage sensor domain (VSD) – which shifts the voltage dependence to more negative voltages– and the other in the pore domain (PD) – which increases the maximal conductance of the channels (Gmax). However, the mechanism by which PUFAs increase the Gmax of the IKs channels is still poorly understood. In addition, it is unclear why IKs channels have a very small single channel conductance and a low open probability or whether PUFAs affect any of these properties of IKs channels. Our results suggest that the selectivity filter in KCNQ1 is normally unstable, contributing to the low open probability, and that the PUFA-induced increase in Gmax is caused by a stabilization of the selectivity filter in an open-conductive state.

## INTRODUCTION

The voltage-gated K^+^ channel KCNQ1, also referred to as Kv7.1, is expressed in the heart. Here the channel associates with the accessory KCNE1 subunit, generating the *so-called* slow delayed-rectifier (IKs) current, an important contributor to the repolarizing phase of the ventricular action potential (AP) ^1,2,3^. Loss-of-function mutations of the KCNQ1/KCNE1 complex are associated with Long QT Syndrome (LQTS) ^4, 5^. These LQTS mutations cause a reduction of the channel current, leading to a significant prolongation of the ventricular AP waveform that can be seen on the electrocardiogram as a prolonged QT interval ^3^ (Fig. 1A). These channel mutations, or dysfunction, increase the risk of developing cardiac arrythmias, which can lead to sudden cardiac death ^6^. At present, treatment of LQTS is mainly based on the usage of β-blockers (most used are long-lasting preparations such as nadolol and atenolol) and implantable cardioverter-defibrillators, especially for patients with high risk of sudden death and frequent syncope ^7^. However, both approaches do not shorten the QT interval duration. Restoring the physiological duration of the QT interval could be achieved by increasing the activity of the KCNQ1/KCNE1 channel complex ^8^ (Fig. 1A).

**Figure 1.**
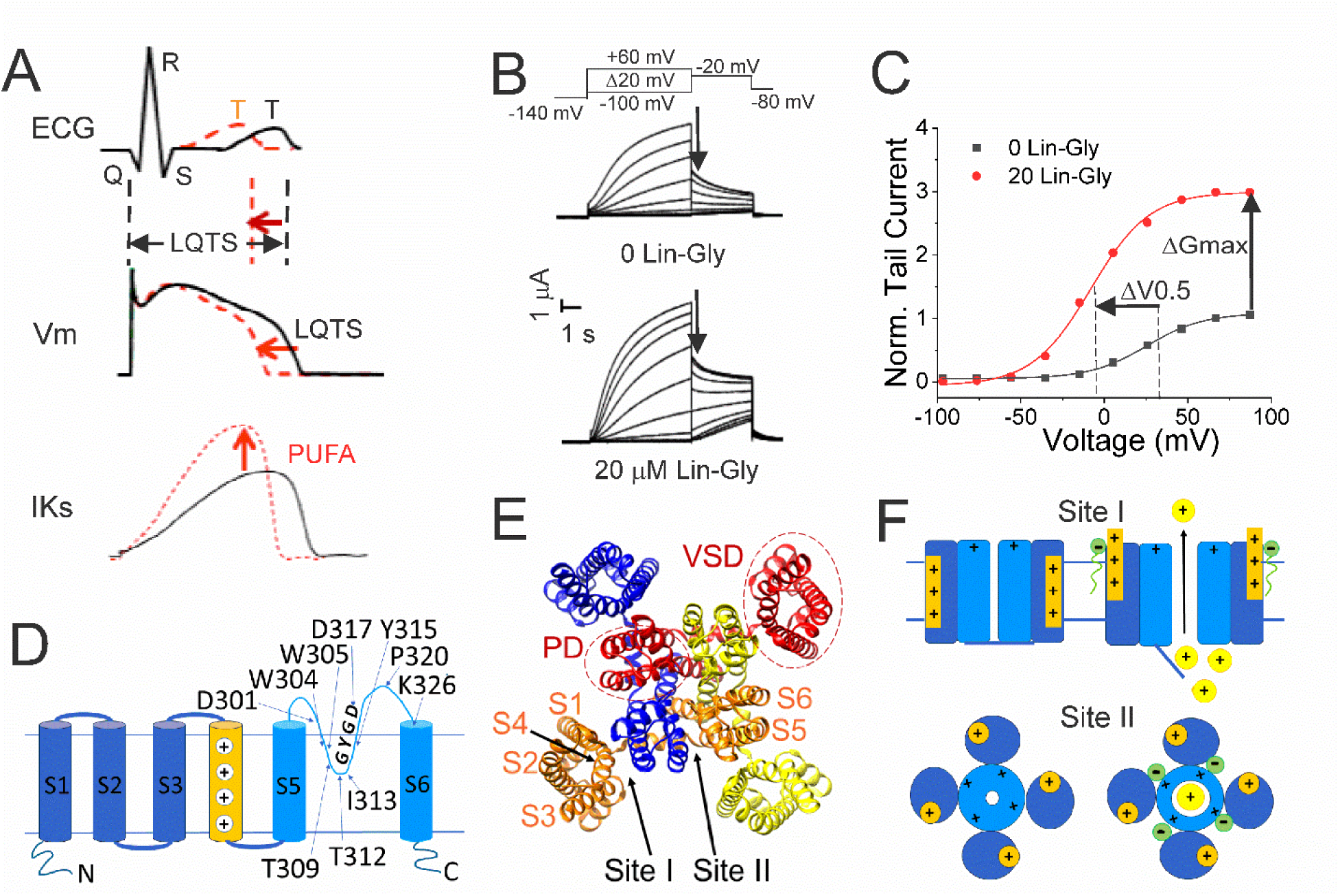
PUFAs activate KCNQ1/KCNE1 channels. A) (black) Prolonged QT interval in the ECG is due to for example loss-of-function mutations of KCNQ1/KCNE1 channels that generate the IKs current that normally contributes to the repolarizing phase of the ventricular AP. (red) PUFAs are potent activators of KCNQ1/KCNE1 channels that can restore the normal functioning of the channel and restore the AP duration and the QT interval. B) Representative current traces of KCNQ1/KCNE1 in 0 μM and 20 μM of Lin-Glycine. Voltage protocol on top. C) Conductance versus voltage curves from tail currents (measured at arrows in B). Channel activation by PUFA results in two main effects: a shift of the voltage-dependence of activation (ΔV0.5) and an increase in the channel maximum conductance (ΔGmax). D) KCNQ1 transmembrane topology. Residues mutated in this study are labeled. E) KCNQ1 top view (PDB: 6UZZ) with PUFA binding sites: Site I, at the VSD; and Site II, at the pore domain. The four subunits are shown in four different colors. F) Cartoon of PUFA mechanism of action. Site I, top panel. Electrostatic interactions between PUFA head groups and positively charged residues in S4 facilitate channel activation by stabilizing the outward state of S4. Site II, bottom panel. PUFA interaction with residues in the pore domain facilitates the increase in the maximum channel conductance.

Like other Kv channels, KCNQ1 has a typical tetrameric structure of four α-subunits. Each α-subunit is composed of six transmembrane segments, with S1-S4 forming the voltage sensor domain (VSD) and S5-S6 forming the pore domain (PD) (Figure 1D). However, KCNQ1/KCNE1 channel complex has a very small single channel conductance and a very low open probability compared to other Kv channels ^9^. The mechanisms behind the small conductance and low open probability are not understood.

Polyunsaturated fatty acids (PUFA), in particular omega 3, are known to exert a protective effect on sudden cardiac death and are recommended in the diet at least twice a week ^10^. PUFAs have been shown to increase IKs currents by a dual mechanism of action, characteristically described as a “Lipoelectric mechanism” ^11^. According to this mechanism, PUFAs can increase IKs currents by shifting the voltage-dependence of activation (ΔV0.5) toward negative voltages and increase the maximum conductance of the channel (ΔGmax) ^12, 13, 14, 15, 16^ (Figure 1B-C).

The two PUFA effects (ΔV0.5 and ΔGmax) on KCNQ1 are independent of each other and originate from the binding of PUFA to two different sites, conventionally indicated as Site I and Site II ^14, 18^. Site I is found in the VSD, where the PUFA head group interacts with the positive charges in the S4 segment and causes the channel to open at more negative potentials ^19^. Site II is found in the pore domain, where electrostatic interactions between the PUFA head group and the positively charged residue, K326, facilitate an increase in the maximal conductance of the channel ^18^ (Figure 1E-F).

Although previous studies have shown the importance of K326 for PUFA in increasing Gmax through binding at site II ^18^, the mechanism by which PUFAs increase Gmax is still not understood. In this study we use a combination of two-electrode voltage clamp, single channel recordings, site-directed mutagenesis, and recent cryoEM structures to gain insight into the molecular mechanism underlying the Gmax effect following PUFA binding at site II. We propose a novel mechanism where the selectivity filter of KCNQ1 is normally unstable and the binding of PUFA to site II stabilizes a network of interactions at the selectivity filter, which in turn, leads to a more open and stable conformation of the KCNQ1 pore and thus an increase in channel open probability.

## Material and methods

### Molecular biology

KCNQ1 (UniProt: P51787), KCNE1 (Uniprot: P15382) cRNA were transcribed using the mMessage mMachine T7 kit (Ambion). KCNQ1 was co-expressed with KCNE1 subunit, following a 3:1, weight: weight (Q1:E1) cRNA ratio to make up the KCNQ1/KCNE1 currents. Site-directed mutagenesis was performed using the Quickchange II XL Mutagenesis Kit (QIAGEN Sciences) for mutations in KCNQ1. 50 ng of complementary RNA was injected into defolliculated Xenopus laevis oocytes (Ecocyte Bioscience, Austin; Xenopus 1 Corp, Dexter) for channel expression. After RNA injection, cells were incubated for 24-96 h, in standard ND96 solution (96 mM NaCl, 2 mM KCl, 1 mM MgCl_2_, 1.8 mM CaCl_2_, 5 mM HEPES, 1M sodium pyruvate and penicillin (10,000 units)-streptomycin (10 mg/ml in 0.9% NaCl; pH 7.5) at 16°C before experiments.

### Two-electrode voltage clamp

KCNQ1/KCNE1 currents were recorded using the two-electrode voltage clamp (TEVC) technique. The recording chamber was filled with ND96 solution (96 mM NaCl, 2 mM KCl, 1 mM MgCl_2_, 1.8 mM CaCl_2_, and 5 mM HEPES; pH 7.5). Pipettes were filled with 3M KCl. The polyunsaturated fatty acid (PUFA) Lin-Glycine was purchased from Cayman Chemicals (Ann Arbor, MI), kept in stock of 100 mM with ethanol at - 20*°*C and diluted in ND96 solution the day of the experiments. Electrophysiological recordings were acquired using Clampex 10.7 software (Axon, pClamp, Molecular Devices). Lin-Glycine was perfused into the recording chamber in a pre-application step until reaching steady state, followed by an (I-V) protocol to measure the current-voltage relationship before and after perfusion. During PUFA application, cells were held at -80 mV and depolarization from -80 mV to 0 mV (5-s) was applied every 30 s, before stepping to -40 mV and then back to – 80 mV. In the I-V protocol, a hyperpolarized step from -80 mV to - 140 mV is applied before 20-mV voltage steps from -100 mV to +60 mV. Tail currents are recorded at -20 mV before returning to holding potential.

### Single channel recordings

Single channel currents were recorded from transiently transfected mouse *ltk-*fibroblast cells (LM cells) using 1.5 μL Lipofectamine 2000 (Thermo Fisher Scientific). Cells were transfected with 1.5 μg of pcDNA3 containing a linked KCNE1-KCNQ1 construct ^20^, to ensure fully KCNE1-saturated complexes, in addition to a plasmid containing green fluorescent protein (GFP) to identify transfected cells. Cells were recorded from 24-48 hours after transfection using an Axopatch 200B amplifier, a Digidata 1440A and pClamp10 software (Molecular Devices, San Jose CA, USA). The bath solution contained (in mM): 135 KCl, 1 MgCl_2_, 1 CaCl_2_, 10 HEPES, 10 Dextrose (pH 7.4 with KOH). The pipette solution contained (in mM): 6 NaCl, 129 MES, 1 MgCl_2_, 5 KCl, 1 CaCl_2_, 10 HEPES (pH 7.4 with NaOH). Pipettes were pulled from thick-walled borosilicate glass (Sutter Instruments, Novato, CA, USA), using a linear multistage electrode puller (Sutter Instruments), fire polished and coated with Sylgard (Dow Corning, Midland, MI, USA). Electrode resistance was between 40 and 80 MΩ after polishing. Currents were sampled at 10 kHz, low-pass filtered at 2 kHz at acquisition and subsequently digitally filtered at 200 Hz for presentation and analysis. Data were collected using cell-attached patch configuration to minimize disruption to the patch and avoid rundown problems due to the loss of PIP_2_. Lin-Glycine was solubilized in DMSO and added directly to the bath. Only patches that were largely free of endogenous currents and had few channels, such that there were several blank sweeps to average for use for leak subtraction, were analyzed.

### Data analysis

To measure the conductance vs voltage (G-V) curve, KCNQ1/KCNE1 tail currents were measured at -20 mV and obtained values were plotted against the activation voltages and fitted to a Boltzmann function:

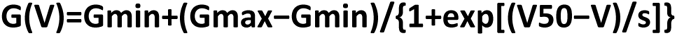

where *G_min_* is the minimal conductance, *G_max_* is the maximal conductance, *V*_50_ the midpoint (which describes the voltage at which the conductance is half the maximal conductance established from the fit), and *s* is the slope of the curve in mV. The difference in Gmax effect, before and after application of Lin-Glycine in each oocyte is used as a measure of the change in maximal conductance. To understand the concentration dependence of LIN-Glycine effect on *G_max_*, the following concentration-response curve was fitted to the data:

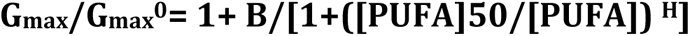

where B is the maximum relative increase in *G_max_* {(*G_max_* - *G_max0_*)/*G_max0_*}, [PUFA]_50_ the PUFA concentration needed to cause 50% of the maximal effect, and H the Hill coefficient. Average values are expressed as mean ± SEM and n represents the number of experiments (unless specified).

Statistical analysis was conducted using GraphPad Prism 8 (GraphPad Software, Boston, Massachusetts, USA). Statistical tests used were One-way ANOVA with Dunnett’s post-hoc multiple comparisons test (Fig. 5) and student t test for all other single comparisons.

Data were analyzed using Clampfit 10.7 (pCLAMP), Origin Pro (OriginLab Corporation) and GraphPad Prism 8 software (GraphPad Software, Boston, Massachusetts, USA).

## RESULTS

### Lin-Glycine drastically increases the chance of KCNQ1/KCNE1 channel opening

Lin-Glycine has been shown to increase the Gmax in whole-oocyte recordings of KCNQ1/KCNE1 channels 2.5-fold ^13^. To better understand how Lin-Glycine increases Gmax, we here extended our analysis to the single-channel level to study the behavior of KCNQ1/KCNE1 in the absence and presence of Lin-Glycine (20 µM) (Fig. 2). Representative traces (Fig. 2A-B) and all points histograms (Fig. 2C-D) suggest that there is no change in single channel conductance between KCNQ1/KCNE1 in control and with Lin-Glycine. This shows that the Gmax effect is not due to an increase in the single channel conductance. However, Lin-Glycine caused a decrease in the latency to first opening (In control, 1.78 ± 0.36 s, n = 41 sweeps and after 20 µM Lin-Glycine, 1.03 ± 0.05 s, n = 99 sweeps; p = 0.0025) and an increase in the number of non-empty sweeps (Suppl. Fig. S1A). In control, there are a high number of empty sweeps as shown before for KCNQ1/KCNE1 channels ^17^. The average current during the single channel sweeps was increased by 2.3 ± 0.33 times (n = 4 patches, p =0.006) by the application of Lin-Glycine (Fig. 2E). The number of non-empty sweeps increased 2.85-fold (2.85 ± 0.85, n = 200 sweeps from 4 patches; p = 0.001), going from a total of 41/200 non-empty sweeps in control to 99/200 when Lin-Glycine was applied. In contrast, once the channel opens, it seems to behave very similarly in control and Lin-Glycine because, in non-empty sweeps, the open probability (Po) in the last second of the traces was almost identical between control and Lin-Glycine conditions (Po = 0.78 ± 0.02 (n = 8 sweeps) in control vs 0.87 ± 0.04 (n = 8 sweeps) in Lin-Glycine) (Fig. 2F). This high open probability was maintained in control conditions even for longer depolarizing voltage pulses, as if once the channels had opened they stayed open for the remaining of the voltage steps (Suppl. Fig. S2). The 2.85-fold increase in the number of non-empty sweeps is very similar to the Gmax increase seen in macroscopic currents.

**Figure 2.**
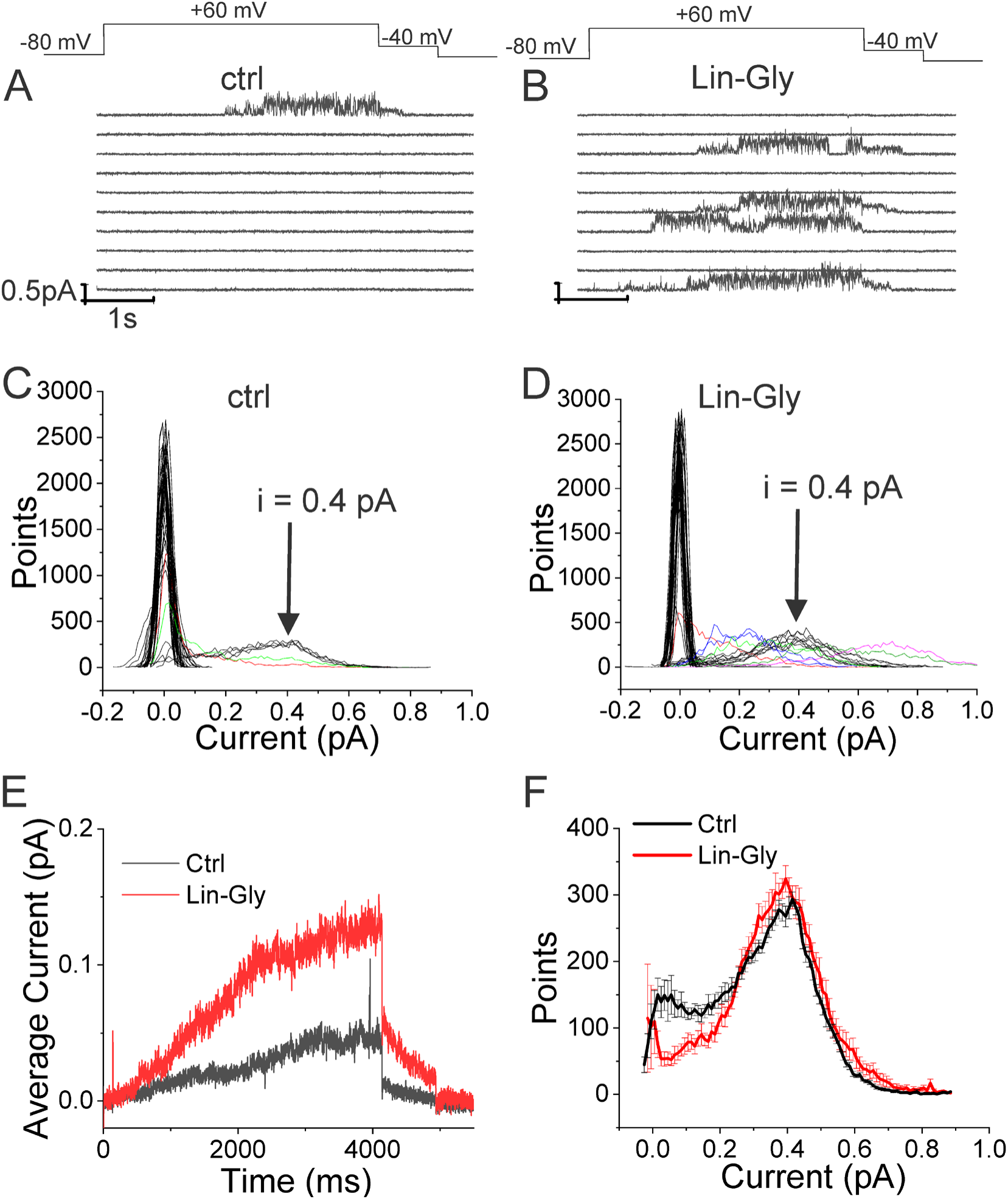
Lin-Glycine induces an increase in the Po of KCNQ1/KCNE1. A-B) 10 consecutive traces of KCNQ1/KCNE1 in A) control and B) in the presence of Lin-Glycine (20 μM) (top) Protocol used for the recordings. C-D. All-point amplitude histogram of 50 consecutive traces in C) control and D) Lin-Glycine). Note no change in the single-channel current amplitude, however an increase in the number of sweeps with channel opening is observed. Note that there were at least two channels in this patch. Different sweeps were assigned different colors to better visualize different types of channel behaviors. Note panels A-D are all from the same patch. E) Average currents of 100 sweeps in control and Lin-Glycine. F) All-point histogram of the last second of non-empty sweeps in control and in Lin-Glycine. We estimated the open probability from the all-point amplitude histogram by Po = Sum (iN/(i_estimate_N_total_)), where N is the number of points for a specific current i in the histogram, i_estimate_ = 0.4 pA from the peak of the histogram, and N_total_ = 10,000 is the total number of points in the last second of the trace. p = 0.78 ± 0.02 (n = 8 sweeps) and p = 0.87 ± 0.04 (n = 8 sweeps) for Control and Lin-Glycine, respectively, from the same patch.

The decrease in first latency is most likely due to an effect of Lin-Glycine on Site I in the VSD and related to the shift in voltage dependence caused by Lin-Glycine. In contrast, the increase in the number of non-empty sweeps is most likely an effect on Site II in the pore and related to the Gmax effect. We conclude that the Gmax effect of Lin-Glycine on KCNQ1/KCNE1 is mainly due to an increase in the Po by increasing the number of non-empty sweeps.

**Supplementary Figure S1.**
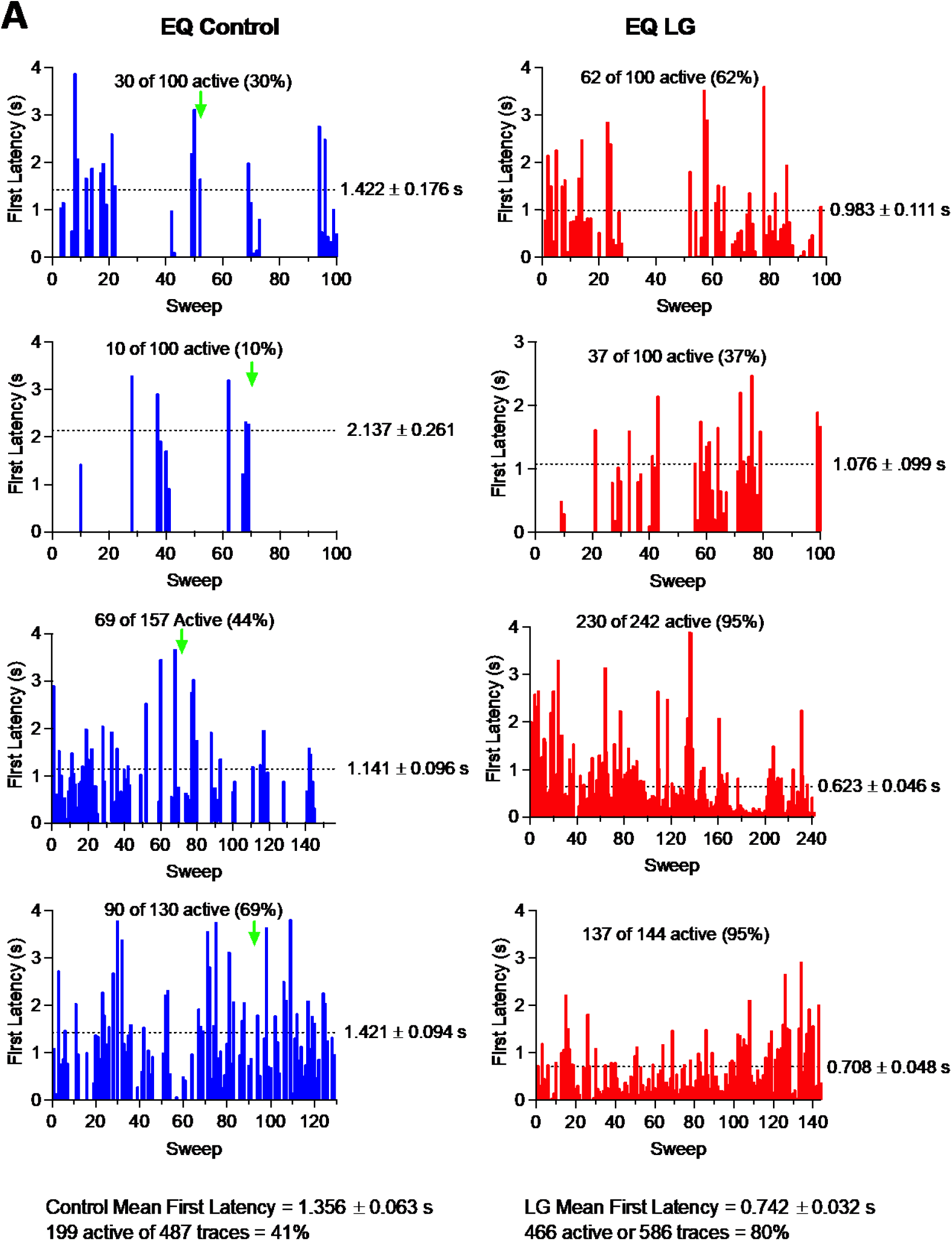
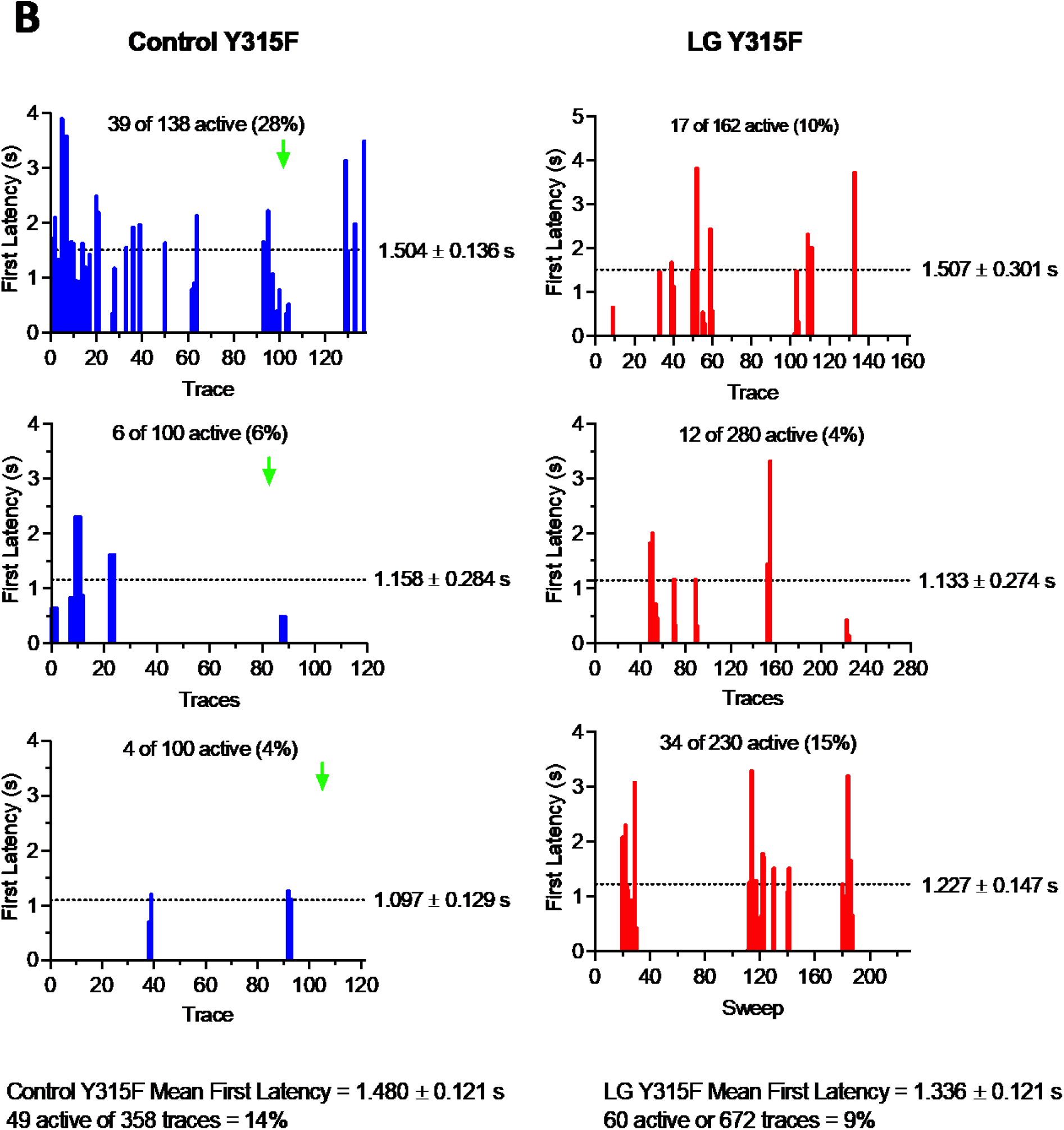
First latency to opening is shortened by Lin-Glycine. A-B) Representative diary plot of first latency to opening during 4 s depolarizations to +60 mV for A) KCNQ1_WT /KCNE1 (EQ) and B) KCNQ1_Y315F/KCNE1 (Y315F) in control (left) and 20 μM Lin-Glycine (right). Latency values of zero correspond to sweeps with no channel openings. Green arrow indicates point at which Lin-Glycine (LG) was added. Mean ± SEM first latency for each cell before and after Lin-Glycine (LG) are indicated next to each plot. Note that these are representative traces and for some patches there are more recordings than shown. In the analysis in the main text we used only traces recorded clearly before application of Lin-Gly and after Lin-Gly had been applied for several minutes, and we used only patches with low number of openings to avoid complications of simultaneous channel openings if more than one ion channel was present in the patch. Note that left and right panels in each row are from the same patch, but each row is from a different patch.

**Supplementary Figure S2.**
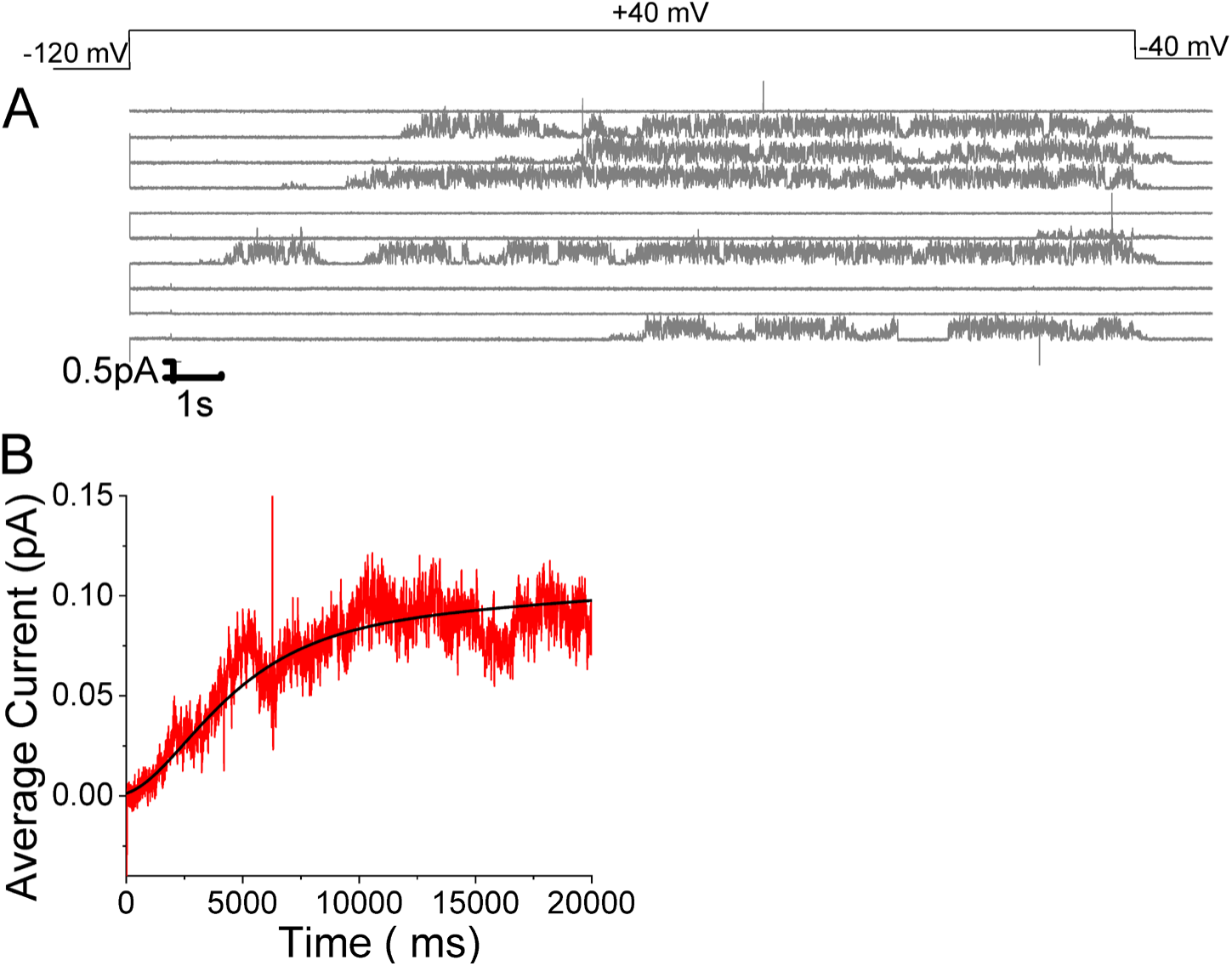
Open probability stays high in KCNQ1/KCNE1 channels once opened. A) 10 consecutive traces of KCNQ1/KCNE1in response to 20-sec long voltage steps in control solutions. (top) Protocol used for the recordings. B) Average currents of 57 sweeps in control solution.

### Crevice residues affect PUFA ability to increase Gmax

It was previously shown that PUFAs binding at Site II electrostatically interact with the positively charged residue K326, located just outside the selectivity filter ^18^. We tested whether another residue very close to K326, the aspartic acid at position 301, is important for the PUFA interaction at Site II. Electrophysiological analysis revealed that when KCNQ1_WT/KCNE1 was mutated to KCNQ1_D301E/KCNE1, the ΔV0.5 was shifted similarly to the WT channel (Supplementary Figure S3) but a dramatic decrease of the Gmax effect of Lin-Glycine was observed, showing the importance of this residue for PUFAs interaction at Site II (Figure 3A-C). However, it is not clear how PUFA binding at Site II increases Gmax.

**Figure 3.**
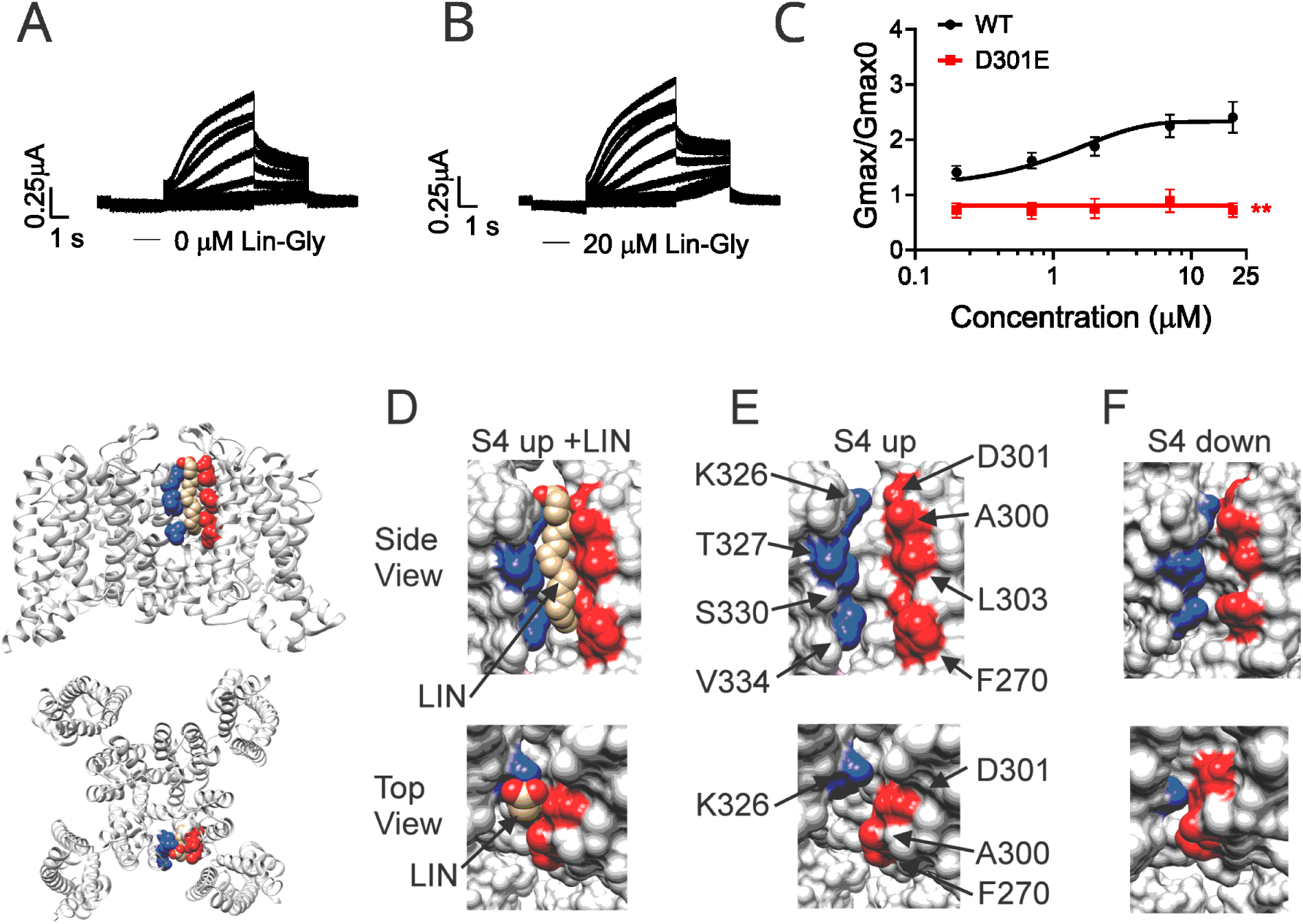
PUFA binds to a state-dependent small crevice between K326 and D301. A-B) KCNQ1_D301E/KCNE1 representative current traces A) in 0 μM of Lin-Glycine. B) After perfusion of 20 μM of Lin-Glycine. C) Gmax/Gmax0 for KCNQ1/KCNE1 channels and KCNQ1_D301E /KCNE1 channels. Gmax/Gmax0 was significantly reduced for the D301E mutation compared to WT channels (p = 0.0018, n = 4 oocytes). Student t test was used to conduct statistical analysis. D, E and F) Structures of crevice between S5 and S6 in KCNQ1 with S4 up (D and E) and S4 down (F). Residues that surround the crevice from S6 shown in blue (K326, T327, S330, V334) and from the pore helix and S5 in red (D301, A300, L303, F270). Remaining KCNQ1 residues shown in purple. On the left is shown the location of the crevice in KCNQ1 (top) side view and (bottom) top view. D) In MD simulations^14^, linoleic acid (LIN: gold color) fits in a narrow crevice present in the cryoEM structure of activated state KCNQ1 (S4 up). E) Same view as, in D but without LIN. F) In the cryoEM structure with S4 in the resting state (S4 down), the crevice between K326 and D301 is too narrow to fit LIN.

In our previous MD simulations ^14^ based on the cryoEM structure of KCNQ1 with S4 in the activated state, PUFA binds to residues K326 and D301 at Site II that delimit a narrow crevice (Figure 3D-E), where PUFA could fit with both the head and the tail (Figure 3D-E). However, in the recent cryoEM structure of KCNQ1 with S4 in the resting state ^21^ the crevice between K326 and D301 is now so narrow that PUFA seems unable to fit into it (Figure 3F). In addition, there are large rearrangements in the selectivity filter when comparing the cryoEM structures with S4 activated (PDB: 8SIK) and S4 resting (PDB: 8SIN) (Figure 4). We have made a video showing the changes and the reorganization of the pore between these two structures (Supplement Video S1). The conformation of the pore significantly varies between what seems like a conductive (PDB: 8SIK) and a non-conductive (PDB: 8SIN) state of the channel. For example, in the structure with S4 in the resting state, Y315 and D317 have swung outwards and left their positions in which they made hydrogen bonds with two tryptophans, W304 and W305, respectively. Y315 belongs to the K^+^ channels signature sequence, a stretch of 8 amino acids including TXXTXGYG, highly conserved among K^+^ channels of different families and D317 is just next to this sequence ^22, 23^. W304 and W305 belong to the aromatic ring cuff (Figure 4C-D) that has been suggested to be important for the stability of the open state of the selectivity filter in Kv channels ^24,25^.

**Figure 4.**
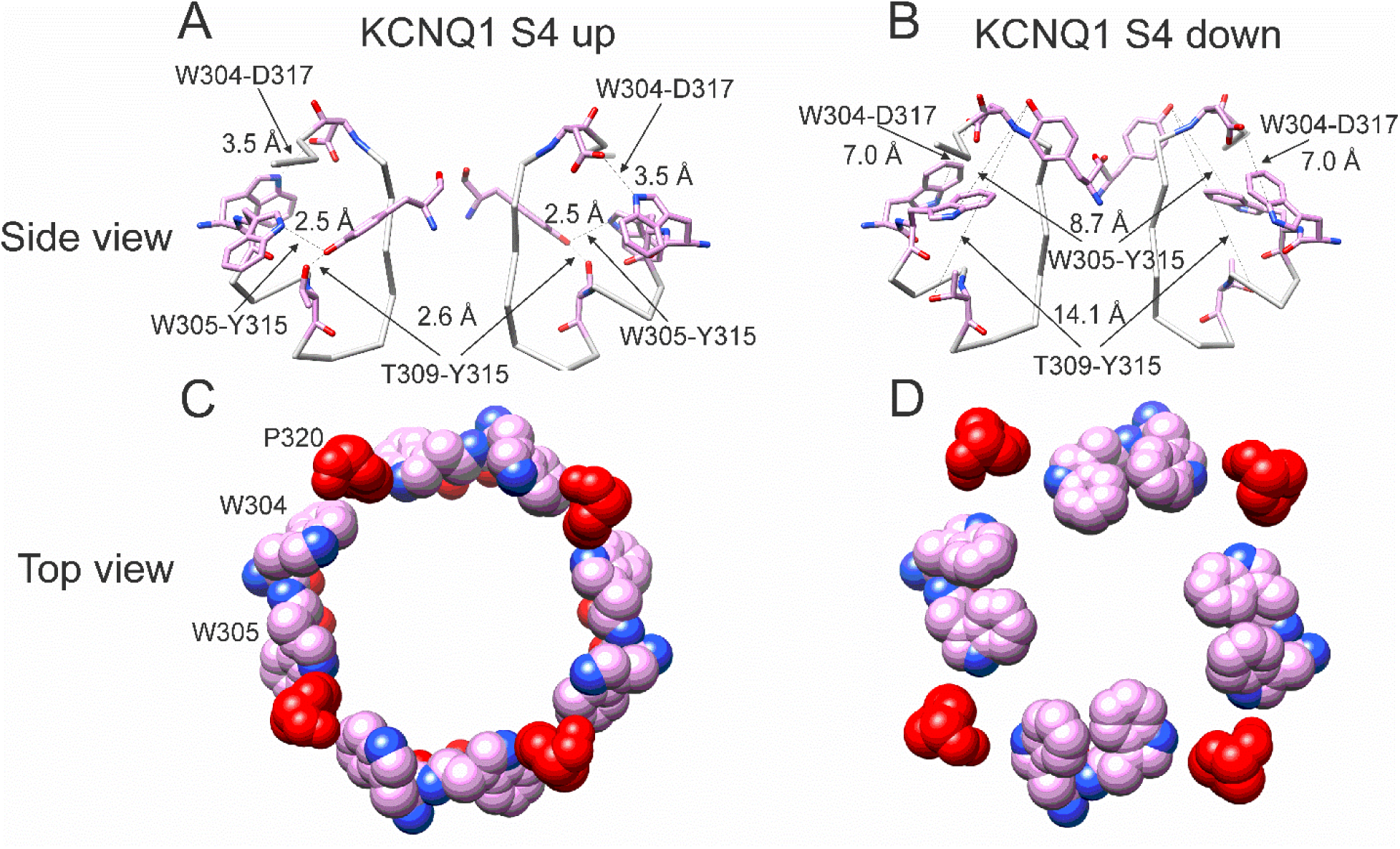
Different conformations of selectivity filter in cryoEM structures with S4 activated or resting. A) Selectivity filter of KCNQ1 with S4 activated (S4 up; PDB: 8SIK). Distances between D317 and W305 and between T309 and Y315 are short enough to form hydrogen bonds (dashed lines). B) Selectivity filter of KCNQ1 with S4 in resting state (S4 down; PDB: 8SIN). Distances between D317 and W305 and between T309 and Y315 are too long for hydrogen bonds (dashed lines). Only two subunits are shown for clarity. C-D). Aromatic cuff in KCNQ1 with C) S4 activated and D) resting S4. Note how P320 (red) moves away from its position in between W304 and W305 from two different subunits in the S4 down conformation.

In addition, P320, which in the structure with an activated S4 sits in between W304 and W305 from two subunits (Figure 4C), has also swung outwards in the structure with S4 in the resting state and exposed a gap between W304 and W305 in the aromatic cuff (Figure 4D, see also supplementary Video S2). The homologous proline in other Kv channels sits between the two tryptophans and stabilizes the aromatic ring cuff. Supplementary Figure S4 and S5 show the extent of the movement for each residue in the selectivity filter and nearby regions between the two conformations.

**Supplementary Figure S3.**
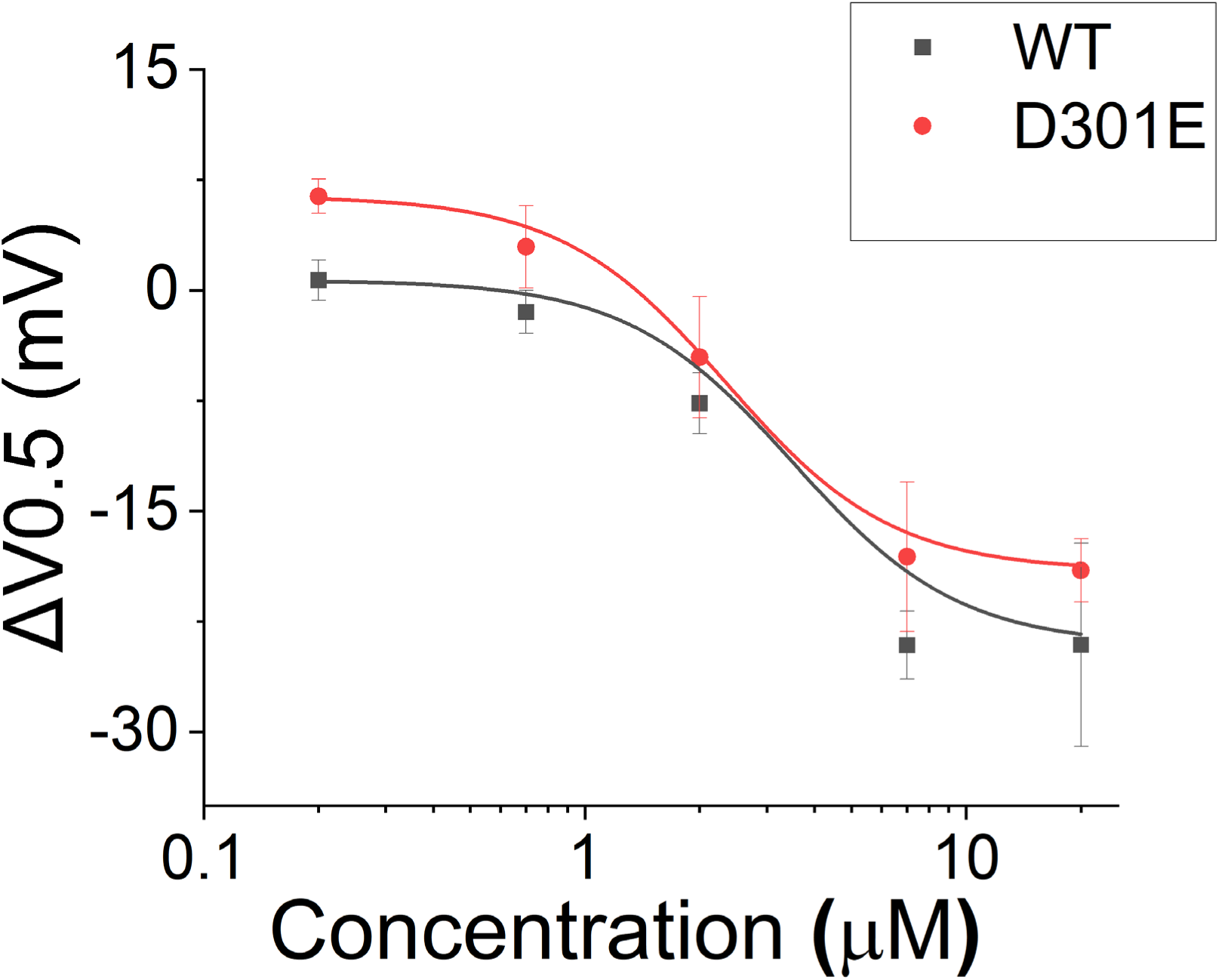
Effect of Lin-Glycine in shifting the voltage-dependence of activation (ΔV0.5). A similar shift in the voltage dependence of activation was found for KCNQ1_D301E/KCNE1 and KCNQ1/KCNE1 after perfusion of several concentrations of Lin-Glycine. Comparisons at 20 μM of Lin-Glycine revealed no significant difference between the effect seen in KCNQ1/KCNE1 and KCNQ1_D301E (p = 0.52; n = 3 oocytes).

**Supplementary Figure S4.**
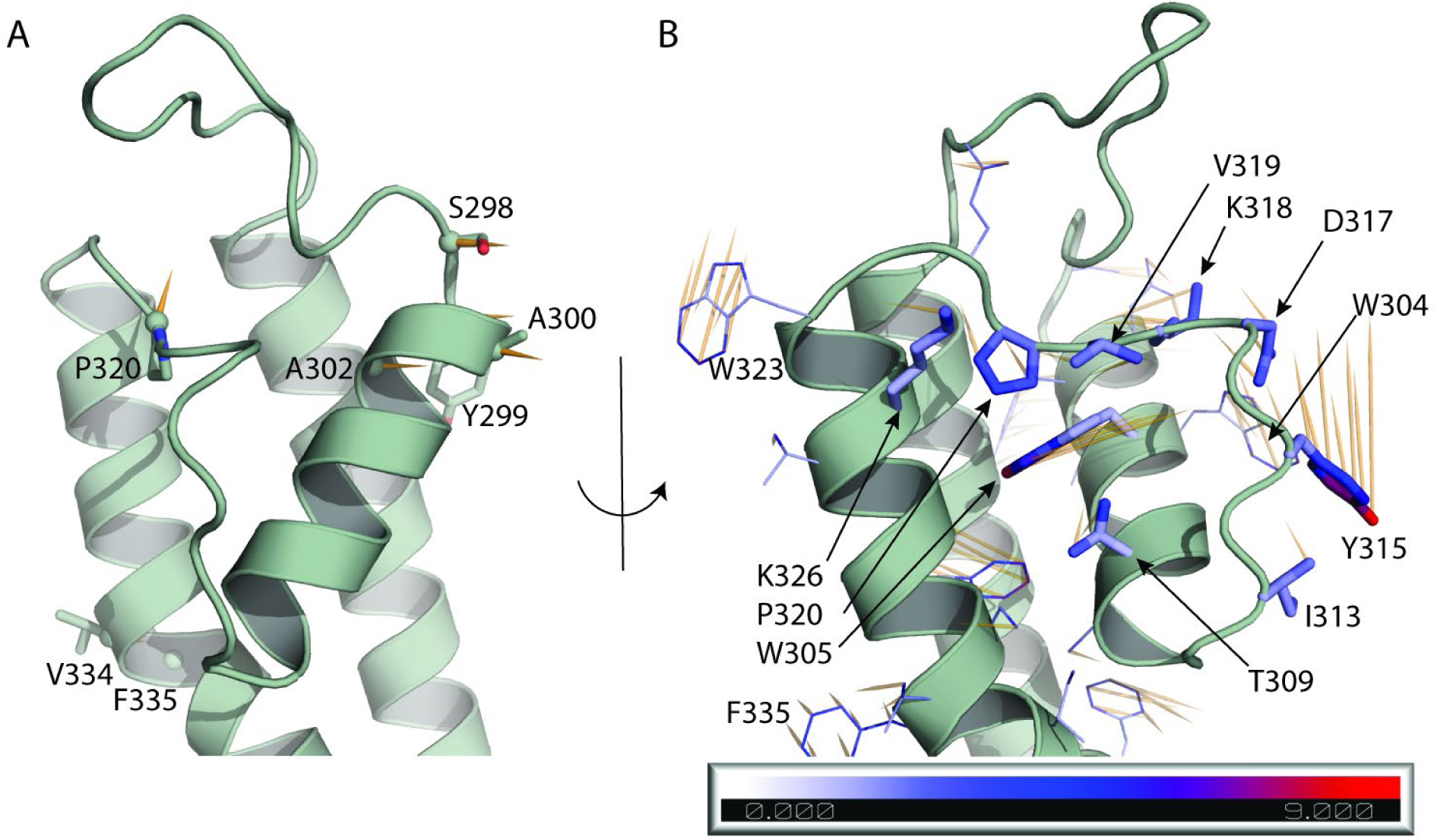
Atomic displacements at the selectivity filter and nearby regions in the transition from up to down state. Each vector (shown in orange) starts from the position of a given atom in the up state (PDB ID 8SIK) and points to the position of the same atom in the down state (PDB ID 8SIN). **A.** Displacement of the Ca atoms of residues 298 to 341 is shown in orange if greater than 2 Å. The side chain of such residues is shown as sticks. **B.** Displacement of the side chains of residues 298 to 341. The side chain of residues W304, W305, T309, I313, Y315, D317, K318, V319, P320, K326 are highlighted as sticks and atoms are colored based on the atomic displacement values. Scale bar is in Å. In A and B, for clarity, only one monomer of the up state of KCNQ1 is shown.

**Supplementary Figure S5.**
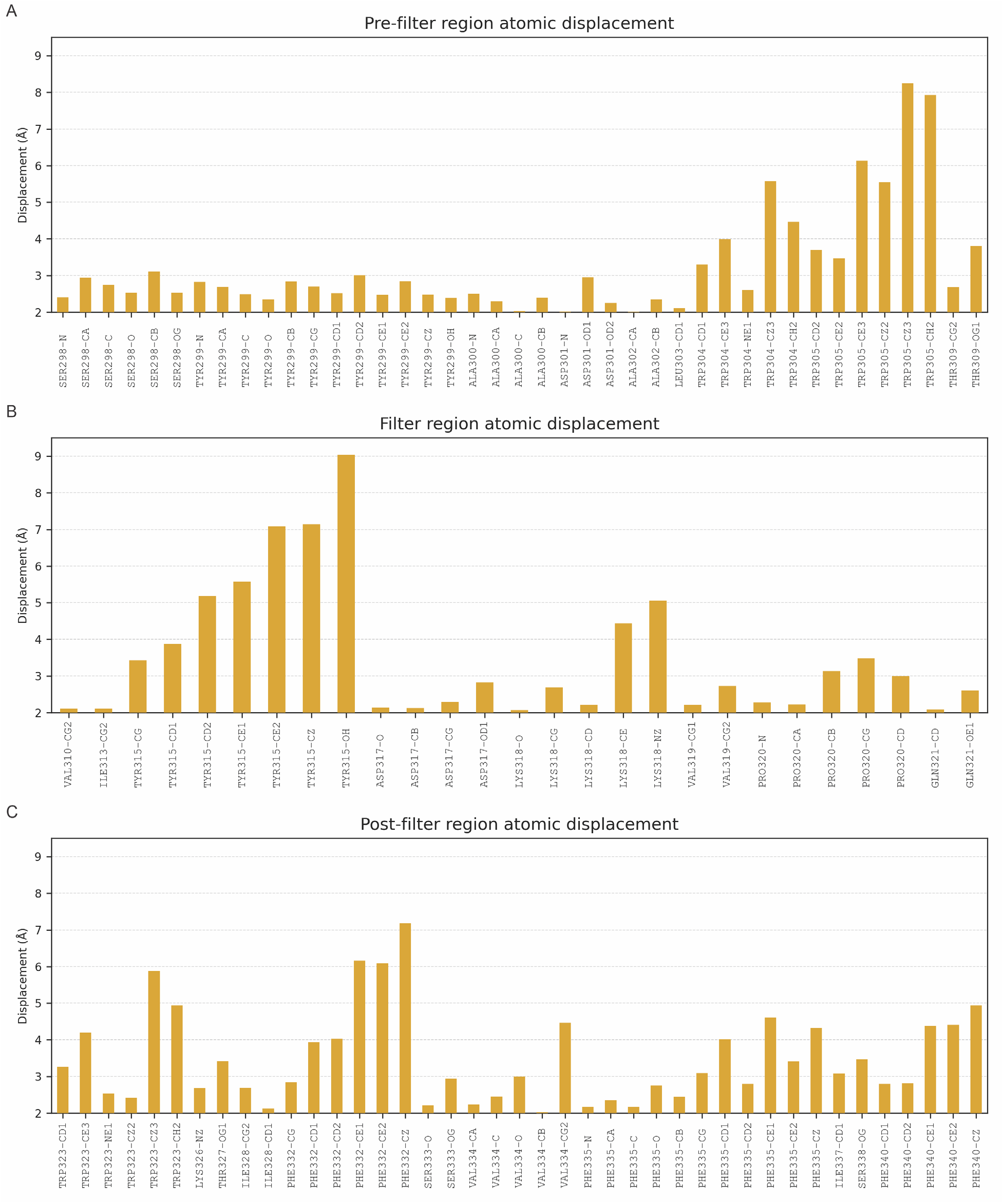
Graphic indicating the atomic displacement of residues at the selectivity filter and nearby regions in the transition from up to down state. Displacement of the Ca atoms of residues 298 to 341 is shown if greater than 2 Å. A) Displacement of atoms (Å) before the selectivity filter; B) residues of the selectivity filter; C) residues located just outside the selectivity filter region,

Based on our single channel data with many empty sweeps, we propose the hypothesis that the KCNQ1 selectivity filter is inherently unstable and transitions between the two conformations - one conductive state that generates the non-empty sweeps and one non-conductive state that generates the empty sweeps in the single channel data. We further propose that PUFA binding to residues at Site II stabilizes and promotes the conductive state of the pore. If PUFA binding to K326 and D301 promotes a series of interactions that stabilize the pore, then residues at the upper part of the selectivity filter, such as Y315 and D317, might be involved in those interactions that promote an open-conductive conformation of the channel.

### Residues that stabilize the selectivity filter are necessary for the Gmax effect

If our pore stability hypothesis is correct, we should see altered effects of Lin-Glycine on Gmax when residues important for the PUFA-promoted open-conductive conformation are mutated. We therefore mutated Y315F and D317E in KCNQ1. We tested Lin-Glycine on KCNQ1_Y315F/KCNE1 and KCNQ1_D317E/KCNE1 channels and compared the Gmax effect to that of the WT channel. The effect of Lin-Glycine was reduced on both mutant channels, with a particularly dramatic decrease for Y315F (Figure 5A). For KCNQ1_Y315F/KCNE1 we compared the effect on the voltage-dependence of activation (ΔV0.5) and we observed a similar effect as for the WT channel (Supplementary Figure S6), with both showing an ∼-25 mV shift. We also tested the Y315F mutation at the single channel level. As expected, Lin-Glycine did not increase the number of non-empty sweeps in KCNQ1_Y315F/KCNE1 channels (52/478 (10.9 % from 3 patches) of traces were non-empty in control and 44/533 (8.3% from 3 patches) of traces were non-empty in Lin-Gly) (Fig. 6 and Supplementary Figure 1B). The mutation Y315F reduced the single channel current slightly from I = 0.4 pA in wildtype to I = 0.3 pA in the Y315F mutation (Fig. 6 C-D). The current average of all traces showed no increase in average current by Lin-Glycine (Fig. 6 E). This data shows that Y315 and D317E are necessary for the ability of Lin-Glycine to increase Gmax. Conductance vs voltage curves (G-V) for WT and mutant channels are shown in Supplementary Figure S7.

**Figure 5.**
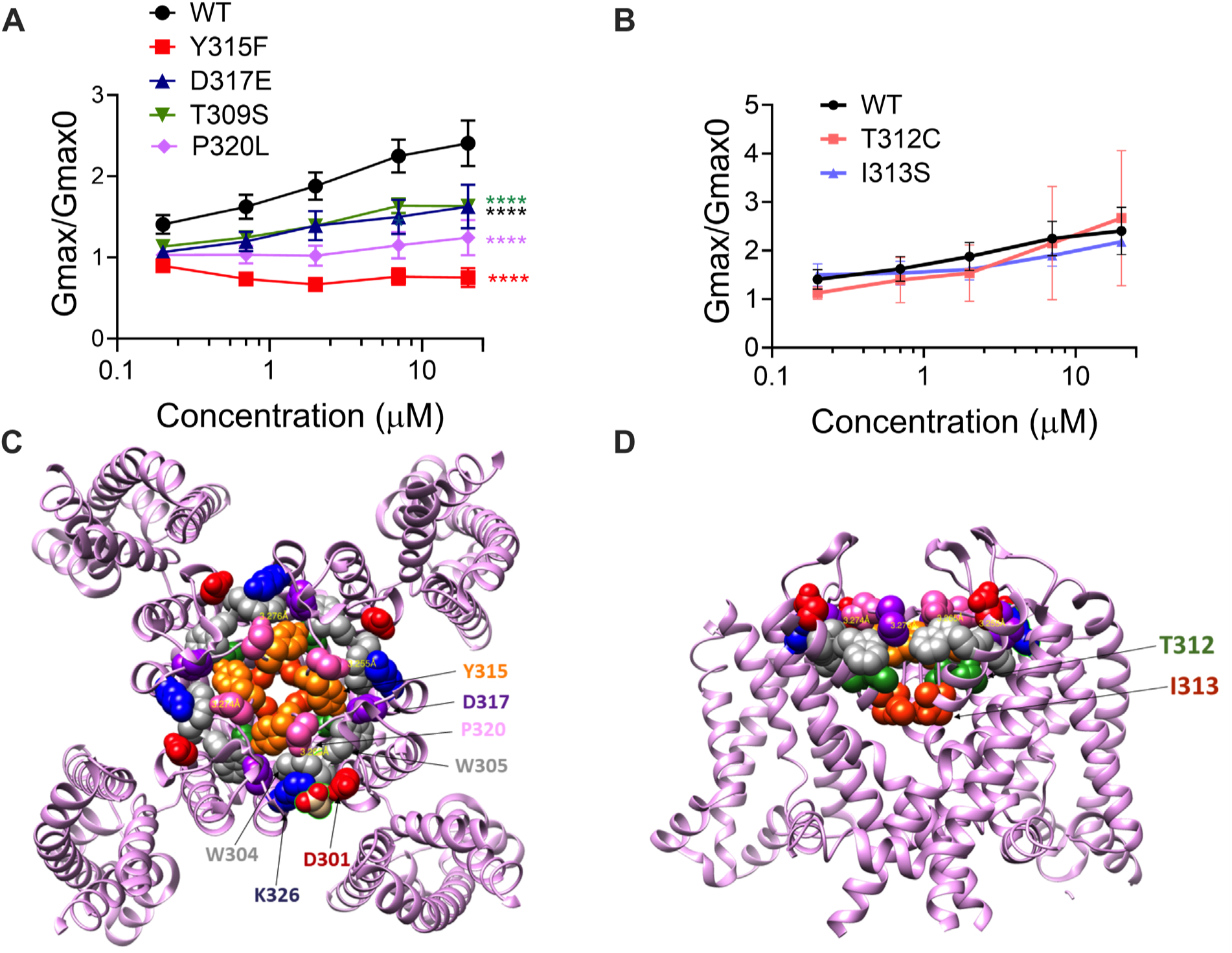
The ability of Lin-Glycine to increase the channel conductance is reduced when channel pore residues are mutated. A) Gmax/Gmax0 values obtained for KCNQ1_WT/KCNE1 channel (black) and mutant channels. KCNQ1_Y315F/KCNE1 (red), KCNQ1_D317D/KCNE1 (blue), KCNQ1_T309S/KCNE1 (green) and KCNQ1_P320L/KCNE1 (purple), the Gmax/Gmax0 is significantly reduced. (p<0.0001(n = 4 oocytes). One-way ANOVA with Dunnett’s post-hoc multiple comparisons test was used for statistical analysis. B) Gmax/Gmax0 values for KCNQ1_T312C/KCNE1 and KCNQ1_I313S/KCNE1 (p=0.14 and p=0.10 respectively. (n= 3 oocytes). One-way ANOVA with Dunnett’s post-hoc multiple comparisons test was used for statistical analysis. C-D) Top view and side view of KCNQ1 channel with mutated residues highlighted.

**Figure 6.**
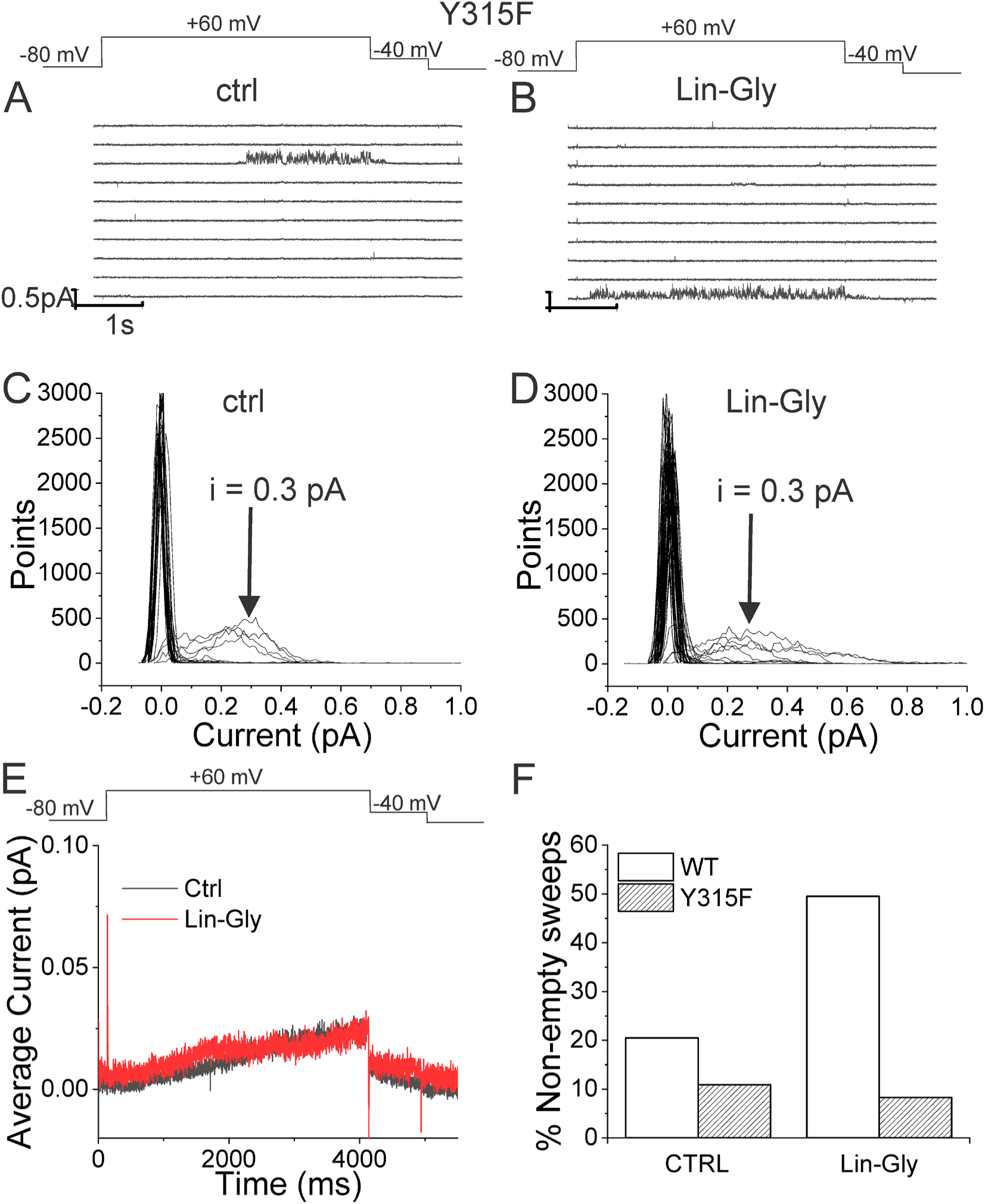
Lin-Glycine does not increase the Po of KCNQ1_Y315F/KCNE1. A-B) 10 consecutive traces of KCNQ1_Y315F/KCNE1 in A) control and B) in the presence of 20 μM Lin-Glycine. (top) Protocol used for the recordings. C-D) All-point amplitude histogram of 50 consecutive traces in C) control and D) Lin-Glycine). The single-channel current amplitude was reduced to 0.3 pA compared to 0.4 pA for WT KCNQ1/KCNE1 (cf. Figure 2C-D). Note that there were at least two channels in this patch. E) Average currents of 478 sweeps in control and 533 sweeps in Lin-Glycine. Note that panels A-D were all from the same patch.

We also found that another residue in the pore helix, T309, was important for the Gmax effect of Lin-Glycine. The homologous residue in the Shaker channel was earlier suggested to be important for the stabilization of the selectivity filter by hydrogen bonding to one of the tryptophans in the aromatic ring ^26^. We generated the mutant channel KCNQ1_T309S/KCNE1 and measured the effect of Lin-Glycine. As shown in Figure 5, the effect of Lin-Glycine on Gmax of the KCNQ1/KCNE1 mutant channel was noticeably reduced compared to the WT channel showing that this residue contributes to the Gmax effect (Figure 5A). We also tested the involvement of proline, which makes up a part of the aromatic cuff in the activated state of the channel (Fig. 4C) by creating the mutant channel KCNQ1_P320L/KCNE1 and found a significant reduction of the Gmax effect for this mutant (Figure 5A). All these data suggest that mutations of residues at the outer portion of the selectivity filter do affect the Gmax increase by Lin-Glycine. These data are consistent with our hypothesis that PUFAs increase the Gmax by affecting interactions that stabilize the selectivity filter.

The residues involved in the Gmax effect are found near the external region of the selectivity filter, suggesting that the network of interactions that are altered during the switch from non-conductive to conductive state are confined to the outer region of the selectivity filter and pore helix of KCNQ1/KCNE1 channels. To test the specificity of the network localization, we mutated two residues, T312 and I313, in the internal portion of the selectivity filter. As a confirmation of our hypothesis, we found that for both KCNQ1/KCNE1 mutant channels, T312S and I313S, the effect of Lin-Glycine in increasing Gmax resembled values obtained in the WT channels (2.49 ± 0.98 and 2.18 ± 0.05, respectively) (Figure 5B). Mutations of residues in the more intracellular region of the selectivity filter do not affect the Gmax increases, as if the interactions that stabilize the channel involve residues located near the external region part of the selectivity filter.

**Supplementary Figure S6.**
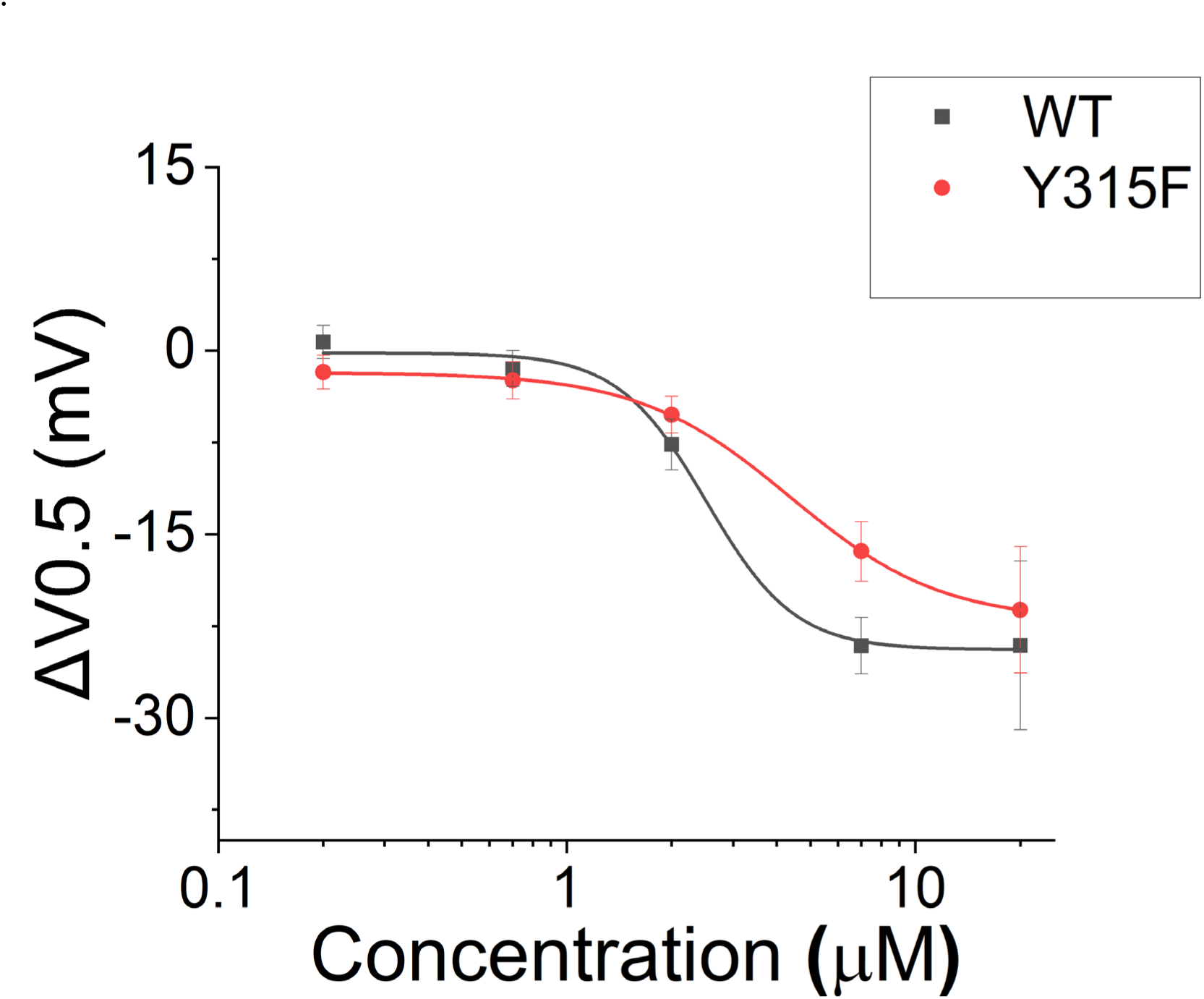
Effect of Lin-Glycine in shifting the voltage-dependence of activation (ΔV0.5). A similar effect in shifting the voltage-dependence of activation was found for KCNQ1_Y315F/KCNE1 and KCNQ1/KCNE1 after perfusion of several concentrations of Lin-Glycine. Comparisons at 20 μM of Lin-Glycine revealed no significant difference between the effect seen in KCNQ1/KCNE1 and KCNQ1_Y315F (p = 0.7435; n = 4 oocytes).

**Supplementary Figure S7.**
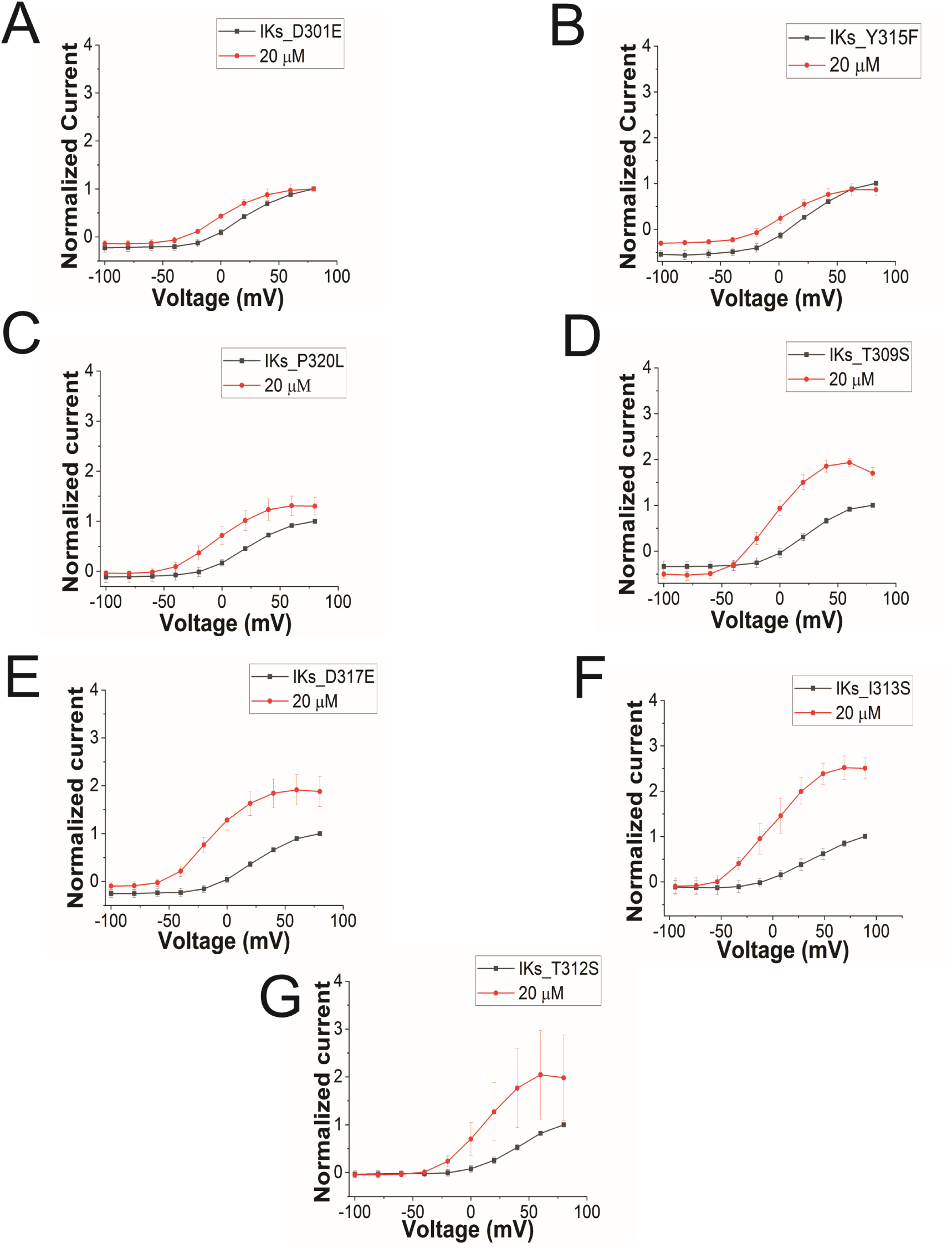
GV curves for KCNQ1 mutants. GV curves in control (black) and after perfusion of 20 μM of Lin-Gly (red) for A) KCNQ1_D301E/KCNE1, B) KCNQ1_Y315F/KCNE1, C) KCNQ1_P320L/KCNE1, D) KCNQ1_T309S/KCNE1, E) KCNQ1_D317E/KCNE1, F) KCNQ1_I313S/KCNE1, and G) KCNQ1_T312S/KCNE1.

Taken together, our results suggest that the binding of PUFA to Site II increases Gmax by promoting a series of interactions that stabilize the channel pore in the conductive state. For instance, we speculate that in the conductive state, hydrogen bonds between W304-D317 and W305-Y315, which are likely absent in the non-conductive conformation of KCNQ1, are created and that PUFA binding to K326 and D301 at Site II favors the transition towards the conductive state of the channel (Figure 7).

**Figure 7.**
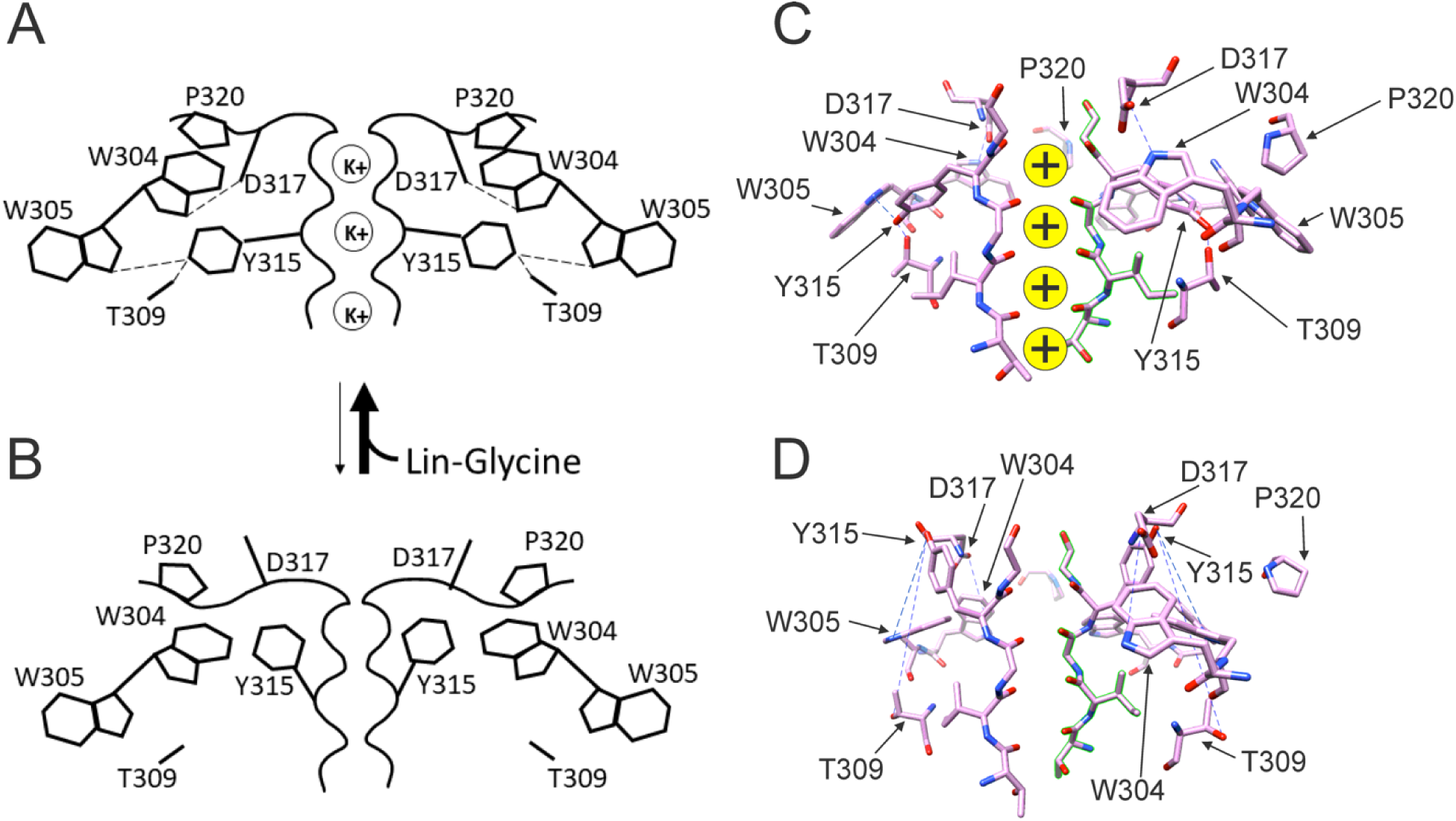
Conformational changes occurring at the pore during the transitions between non-conductive and conductive states. A) The binding of PUFA to site II between K326 and D301 induces a series of interactions between residues near the external part of the selectivity filter. W304-D317 and W305-Y315 form a hydrogen bond (dash line, black). Furthermore, Y315 interacts also with T309 (dash line, black). P320 is reoriented to sit on top of W304 and W305 to favor a more stable configuration of the aromatic ring cuff. The result of those new interactions is a more stable and conductive pore. B) In the non-conductive state those interactions are likely to be absent and this results in a more unstable selectivity filter. Also, P320 is now flipped from its position on top of the two tryptophan of the aromatic ring cuff. C) Cryo-EM selectivity filter with S4 in the activated-state representative of a conductive selectivity filter. D) Cryo-EM selectivity filter with S4 in the resting state, representative of a non-conductive selectivity filter.

## DISCUSSION

We have previously proposed models in which the effect of PUFAs on IKs channels involve the binding of PUFAs to two independent sites: one at the voltage sensor (Site I) and one at the pore domain (Site II). The result is a potent activation of IKs channels, with an increase of the maximum conductance by binding to Site II and a shift of the voltage dependence of activation of the channel by binding to Site I. The mechanism of the voltage shift effects at Site I, where the PUFA head group electrostatically interacts with positively charged S4 residues and thereby facilitates channel activation, has been investigated in previous studies^14, 27, 18, 28^. However, less is known about the molecular mechanism by which PUFA increases the maximum conductance (Gmax) of the channel following binding to Site II. A positively charged lysine at position 326 was suggested to be critical for PUFA-channel interaction, as mutations of K326 completely abolished the Gmax effect ^18^. We have here shown that the interaction at Site II also involves another residue, the aspartic acid 301, as mutagenesis analysis revealed that the effect of Lin-Glycine on Gmax was abolished when the residue was substituted with a glutamic acid.

Our single channel recordings revealed that Lin-Glycine decreases the latency to first opening and increases the Po. The decrease in the latency to first opening is most likely due to the binding of Lin-Glycine to Site I at the VSD and related to the shift in voltage dependence caused by Lin-Glycine (Suppl. Fig S1). In addition, our single channel recordings revealed that Lin-Glycine did not change the single channel conductance, but instead increased Gmax by increasing the Po of the channel. This increase in Po was mainly due to an increase in non-empty sweeps when Lin-Glycine was applied in comparison to control solutions.

We tested the hypothesis that the effect of Lin-Glycine involved conformational changes in the selectivity filter following PUFA binding to two residues K326 and D301 at the pore domain. Those residues delimit a small crevice that seems to change in size in different structures with S4 up or down (Figure 3, D-F). Those residues seem to affect PUFA ability to increase Gmax. We made several mutations of residues known to stabilize the selectivity filter in potassium channels (Y315F, D317E, T309S and P320L; Fig. 5A). The Lin-Glycine effect on the Gmax was much reduced in these mutants suggesting that these residues are necessary for the Gmax effect. To gain further insight into the molecular interactions that could underline the Gmax effect by PUFA binding to site II, we used the latest KCNQ1 structure with the S4 segment in the resting state ^21^. The selectivity filter in this structure showed a very different conformation of the pore region and selectivity filter of KCNQ1 compared to the structures with activated VSDs. Clearly, there will be other differences in the pore domain between structures with activated and resting VSDs, for example the state of the activation gate. Many of the interactions that have been shown to be important for the stability of the selectivity filter are missing in the KCNQ1 structure with a resting VSD. These changes in the selectivity filter can best be seen in our interpolation video between the states with the S4 segments moving from the resting state to the activated state (Supplement video 1). This gave us the idea that the effect of PUFAs in increasing the maximum conductance of the KCNQ1 channel is linked to their ability to stabilize the pore of the channel in a conductive state. It was previously shown that several interactions at the pore region of K^+^ channels are important for ensuring channel conductivity. For example, a feature conserved among K^+^ channels is the aromatic ring cuff that stabilizes the conducting state and plays a role in C-type inactivation ^24, 25, 26, 29^. This structure is made up of two tryptophan residues (W) and a proline residue (P) that sits in between the two tryptophan residues (Supplement video 2). In Shaker and KcsA K^+^ channels, the two tryptophans are involved in C-type inactivation and modification of those interactions manipulate the extent of inactivation ^30, 31^. In Shaker, breaking the hydrogen bonds between the two tryptophans of the aromatic ring cuff and the aspartic acid and tyrosine at the selectivity filter causes an acceleration of the rate C-type inactivation ^26^. In KCNQ1, the two tryptophans of the aromatic ring cuff correspond to W304 and W305. Another conserved residue in K^+^ channels important for the stability of the selectivity filter is P320 (equivalent to P450 in Shaker) that makes up part of the aromatic ring cuff. In the transition between the non-conductive and conductive state of KCNQ1, the proline pulls away from its position close to W304 and W305 and flips outwards (Supplement Video 2). We hypothesize that those hydrogen bonds between W304-D317 and W305-Y315 and the proline interactions are likely absent in the non-conductive conformation of KCNQ1, thereby generating the high number of empty sweeps in control conditions. Occasionally the channel will reform these bonds and interactions and transition into the conductive conformation, thereby generating the non-empty sweeps. We hypothesize that the binding of Lin-Glycine to K326 and D301 at site II biases the transition towards the conductive conformation, thereby increasing the number of non-empty sweeps and increasing Gmax. We here show that mutations of residues involved in these hydrogen bonds in the selectivity filter greatly reduce or abolish the Gmax effect of Lin-Glycine, as if Lin-Glycine increases the Gmax by stabilizing a conductive state of the selectivity filter through these hydrogen bonds.

We noticed that the arrangement of the aromatic ring cuff is slightly different between Shaker K^+^ channels and KCNQ1 channel. For instance, in Shaker, the proline residue sits in between the two tryptophan residues and stabilizes the aromatic ring cuff (Supplementary Figure S8A). In contrast, in KCNQ1 the proline is positioned a little further outward and away from the two tryptophan residues, thus generating a looser arrangement of the aromatic ring cuff (Supplementary Figure S8B). This difference might contribute to rendering the pore of KCNQ1 more unstable, resulting in a large number of empty sweeps and the characteristic flickering nature of channel openings ^32^. In Shaker channels the aromatic cuff seems more stable since it displays few empty sweeps and fewer flickering during bursts. This difference in the stability of the aromatic cuff between Shaker and KCNQ1 might explain why the effect of PUFA on Gmax is large for KCNQ1 but not seen in Shaker ^33, 34^ even if the residues involved in the Gmax effect are conserved between these two channels.

**Supplementary figure S8.**
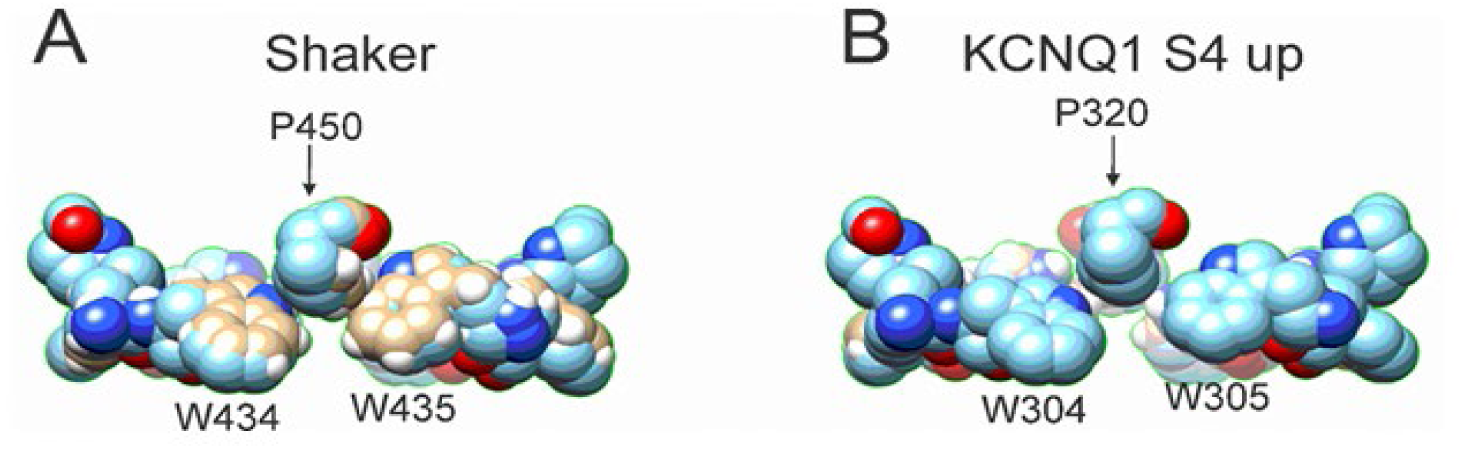
Comparison between the aromatic ring cuff configuration of Shaker and KCNQ1 channels. A) In Shaker, P450 sits in between the two tryptophan and stabilizes the aromatic ring cuff. B) In contrast, in KCNQ1 the P320 is positioned further outward and away from the two tryptophan, generating a looser arrangement of the aromatic ring cuff.

Our single channel data show that the KCNQ1/KCNE1 switches slowly (>10s seconds) between conductive and non-conductive states giving rise to many empty sweeps and a few active sweeps. However, once the channel is turned conductive during a depolarization, the channel stays conductive for the remainder of the sweep (See e.g. Suppl Fig. S2 for 20-s sweeps), as if VSD activation stabilizes the conductive state. Therefore, we propose a model in which the selectivity filter is stabilized more in the non-conductive state when VSD is resting but stabilized more in the conductive state when VSD is activated. This model is consistent with the cryoEM structures of KCNQ1 with VSD resting displaying a non-conductive selectivity filter and with VSD activated displaying a conductive selectivity filter ^21^. The KCNQ1 structure with VSD resting was obtained in a low K^+^ solution, which in other Kv channels would promote C-type inactivation of the selectivity filter. However, we think it is the resting state of the VSD, and not the low K^+^, that caused the non-conductive selectivity filter in the KCNQ1 structure, because a KCNQ1 structure with VSD activated in the low K^+^ solution had a selectivity filter that was nearly identical to the KCNQ1 structure in high K^+^ with VSD activated (except for fewer K^+^ ions in the filter in low K^+^ solutions) (See Suppl. Fig S5 C and D in ^21^). Note also the currents from KCNQ1/KCNE1 channels do not have any external K^+^ dependence (except for what is expected from changes in driving force) and that KCNQ1 channels are inhibited by high extracellular K^+^ concentrations^35^ in contrast to other Kv channels, such as Shaker K^+^ channels, which are inactivated more in low K^+^ ^25, 29^. Our studies suggest that the mechanism of PUFAs to increase KCNQ1/KCNE1 maximum conductance relies on the ability of PUFAs to favor the conductive conformation of the selectivity filter: PUFAs promote interactions between residues in the selectivity filter that stabilize the channel pore. Furthermore, we identify a crevice between K326 and D301 that is present in structures of KCNQ1 with activated VSD but not in structures where the VSD is in the resting state. We propose that PUFA binding in this crevice stabilizes the network of interactions that form the conductive form of the selectivity filter. The result is a more stable and conductive pore, which explains the increase in Gmax by PUFAs.

## Notes

### Competing Interest Statement

The authors have declared no competing interest.

### Summary of Updates

Minor corrections to figures and text and an additional paragraph in the Discussion about how we think the channel functions.

